# Oxygen production via NO dismutation in different ammonia oxidizers

**DOI:** 10.1101/2023.06.07.544047

**Authors:** A. Elisa Hernández-Magaña, Donald E. Canfield, Beate Kraft

## Abstract

Ammonia oxidizing archaea (AOA) are widespread and highly abundant in nature. Despite their typical aerobic metabolism, they can be abundant in ecosystems where oxygen is scarce. Recent observations revealed that the AOA isolate *Nitrosopumilus maritimus* produces oxygen and dinitrogen at nanomolar concentrations, upon oxygen depletion through nitric oxide (NO) dismutation. Here, we explore NO dismutation capability in other ammonia oxidizers with different phylogenetic affinities and from different environmental settings. The organisms explored include three marine AOA, one soil AOA and two soil ammonia-oxidizing bacteria (AOB). Upon oxygen depletion all isolates accumulated oxygen. In incubations with ^15^N tracers with ongoing oxygen accumulation, the AOA strains *Nitrosopumilus adriaticus* and *Nitrosopumilus viennensis* produced ^46^N_2_O from nitrite. Transient ^46^N_2_O accumulation followed by ^30^N_2_ production was detected in the AOA strains *Nitrosopumilus piranensis* and *Nitrosopumilus* sp. CCS1, supporting the earlier observation that NO-dismutation is a common metabolism in AOA, albeit with physiological variations between different strains. An important physiological variable is the capability to reduce N_2_O to N_2_. The finding of oxygen production in several AOA, as well as AOB, indicates that this process is widely distributed among the tree of life and adds an explanation for their abundance in oxygen-depleted environments.

## Introduction

Ammonia-oxidizing archaea (AOA) and ammonia-oxidizing bacteria (AOB) perform the first step in the nitrification process, in which they use oxygen to oxidize ammonia (NH_3_) to nitrite (NO_2-_) (Ward, 2008; Thamdrup, 2012; Kuypers *et al*., 2018). Ammonia oxidizers are widespread in a variety of environments including soils, hot springs, wastewater treatment systems, freshwater and marine ecosystems including sediments and the water column (Alves et al., 2018; Kuypers et al., 2018; Santoro et al., 2019). Most AOB belong to the Betaproteobacteria and Gammaproteobacteria, while AOA belong to the phylum *Nitrososphaerota* (*Thaumarchaeota*) (Alves *et al*., 2018; Santoro *et al*., 2019).

Both AOA and AOB oxidize ammonia to hydroxylamine and further on to nitrite using oxygen. The first step is performed by an ammonia monooxygenase (AMO) (de Bruijn *et al*., 1995; Vajrala *et al*., 2013). The requirement for molecular oxygen to activate AMO has been proven directly in AOB like *Nitrosomonas europaea* through stable isotope assays (Dua *et al*., 1979; Hollocher *et al*., 1981; Klotz *et al*., 1997). Due to the crucial role of oxygen in ammonia oxidation, a decrease in its concentration influences AOB physiology and oxygen concentrations below 25 µmol·l^−1^ can negatively affect AOB growth (French *et al*., 2012). Extensive research on the metabolism of AOB upon the onset of anoxia shows that they can switch to an anaerobic process called nitrifier denitrification, producing NO, N_2_O and in some cases N_2_ (Fochht, Dennis, 1985; Bock *et al*., 1995; reviewed in Stein, 2011).

In the case of AOA, studies on their physiological response to oxygen concentration suggest that they can sustain ammonia oxidation at oxygen concentrations down to around 10µM, (Qin *et al*., 2017). However, below 1µM of dissolved oxygen, ammonia oxidation rates were not detectable and thus AOA were considered inactive (Martens-Habbena *et al*., 2009; Qin *et al*., 2015, 2017, see supplementary information therein).

High AOA abundances have been reported in oxygen-depleted or anoxic ecosystems, such as the Black Sea (Lam *et al*., 2009; Sollai *et al*., 2019), anoxic waters of the Baltic Sea (Berg *et al*., 2015) or in oxygen-minimum zones (OMZs) of the Eastern Tropical South Pacific (ETSP) (Molina *et al*., 2010) and the Eastern Tropical North Pacific (ETNP). In these OMZs, AOA were abundant in depths with oxygen concentrations below the detection limit (1µM) (Pajares *et al*., 2019). The abundance of AOA in these “anoxic” ecosystems is enigmatic and sometimes explained as result of oxygen intrusions or local production of oxygen by phototrophs (Ulloa *et al*., 2012; Garcia-Robledo *et al*., 2017).

The mechanisms by which AOA cope with oxygen limitation or anoxia remained unclear until the recent description of a previously overlooked pathway. In this pathway, upon oxygen depletion, nanomolar concentrations of oxygen and dinitrogen were produced by the AOA *Nitrosopumilus maritimus* SCM1 (Kraft *et al*., 2022). The proposed mechanism is a dismutation of nitric oxide (NO) to O_2_ and N_2_O (Eq.1), which is exergonic (ΔG0’= −165 kJ·mol^−1^) (Kraft *et al*., 2022).

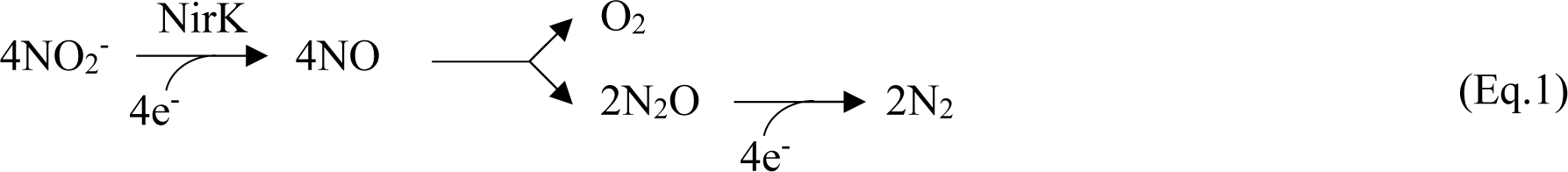

In the proposed pathway of oxygen and (ultimately) dinitrogen production in *N. maritimus* (Eq.1), nitrite is reduced to nitric oxide by the NirK nitrite reductase. Nitric oxide is dismutated into oxygen and nitrous oxide, which is further reduced to dinitrogen.

A similar NO dismutation pathway has been observed previously in the methane-oxidizing bacterium *candidatus* Methylomirabilis oxyfera (NC10), where NO_2-_ is reduced to NO by a NirS nitrite reductase, and two NO molecules are subsequently disproportionated into N_2_ and O_2_. In this pathway, 75% of the intracellularly produced O_2_ is used for methane oxidation, while the rest of the oxygen produced is used for other purposes (Ettwig *et al*., 2010, 2012; Wu *et al*., 2011). In contrast to the archaeaon *N. maritimus,* NC10 produces N_2_ directly in the dismutation step.

So far, the only AOA for which oxygen production has been shown is *N. maritimus* SCM1 (Kraft *et al*., 2022). However, the enzymatic machinery for NO-dismutation remains unidentified, making difficult to explore the potential distribution of the pathway in the environment and its ecological importance. Therefore, the main objective of this study is to explore the NO-dismutation capability in different AOA isolates in pure culture, as well as in other nitrifiers not phylogenetically related to *Nitrosopumilus*. In particular, we also explored the physiology of two AOB strains and their potential to produce oxygen under oxygen depletion. Our approach was to use trace-range oxygen sensors (nanomolar range) and assays with ^15^N labelled substrates.

## Material and Methods

Oxygen production in the dark coupled to dinitrogen production was tested in oxygen-depleted incubations of four marine AOA strains from the clade *Nitrosopumilales* (Group I.1a), *Nitrosopumilus piranensis* NF5, *Nitrosopumilus adriaticus* DC3, isolated from the Adriatic Sea (Bayer *et al*., 2016; Bayer, Vojvoda, *et al*., 2019) and the *Nitrosopumilus* sp. CCS1 a novel strain isolated from the water column of the California Current system in the North Pacific Ocean (Bayer *et al*., 2022). The marine strain *N. maritimus* SCM1, previously tested in Kraft *et al*. (2022), was also included. The soil AOA strain from the clade *Nitrososphaerales* (Group I.1b), *Nitrososphaera viennensis* (Tourna *et al*., 2011) was tested as well. The AOB strains tested were *N. europaea* and *Nitrosospira multiformis*.

The strain *N. maritimus* SCM1 (NCIMB 15022) was provided by Martin Könneke. The strain *N. piranensis* DC3 (JCM 32271, DSM 106147, NCIMB 15115) and *N. viennensis* EN76 (JCM 19564, DSM 26422) were obtained from the Japan Collection of Microorganisms (JCM), the strains *N. adriaticus NF5* (JCM 32270, NCIMB 15114) and *Nitrosopumilus* sp. CCS1 were obtained from the culture collection of Alyson Santoro at the University of California, Santa Barbara, while the strains *N. europaea* (DSM 28437) and *N. multiformis* (DSM 101674) were obtained from the DSMZ-German Collection of Microorganisms and Cell Cultures GmbH.

Axenic batch cultures of the AOA strains *N. piranensis* DC3, *N. adriaticus* NF5 (Bayer *et al*., 2016; Bayer, Vojvoda, *et al*., 2019) and *Nitrosopumilus* sp. CCS1 (Bayer *et al*., 2022) were grown in Synthetic Crenarchaeota Medium (SCM) – HEPES buffered (pH 7.6) at their respective optimal temperature (Table 1) with the addition of catalase (Bayer, Pelikan, *et al*., 2019). *N. viennensis* EN76 was maintained in freshwater medium for soil Archaea (Tourna *et al*., 2011). The ammonia-oxidizing bacteria (AOB) *N. europaea* and *N. multiformis* were grown in axenic batch cultures in basal mineral salt medium (Koops *et al*., 1991) HEPES-buffered (pH 7.6) at 28°C. Medium 1583 for ammonia-oxidizing bacteria (DSMZ, 2007). All the cultures were kept in the darkness and without stirring.

**Table 1.**
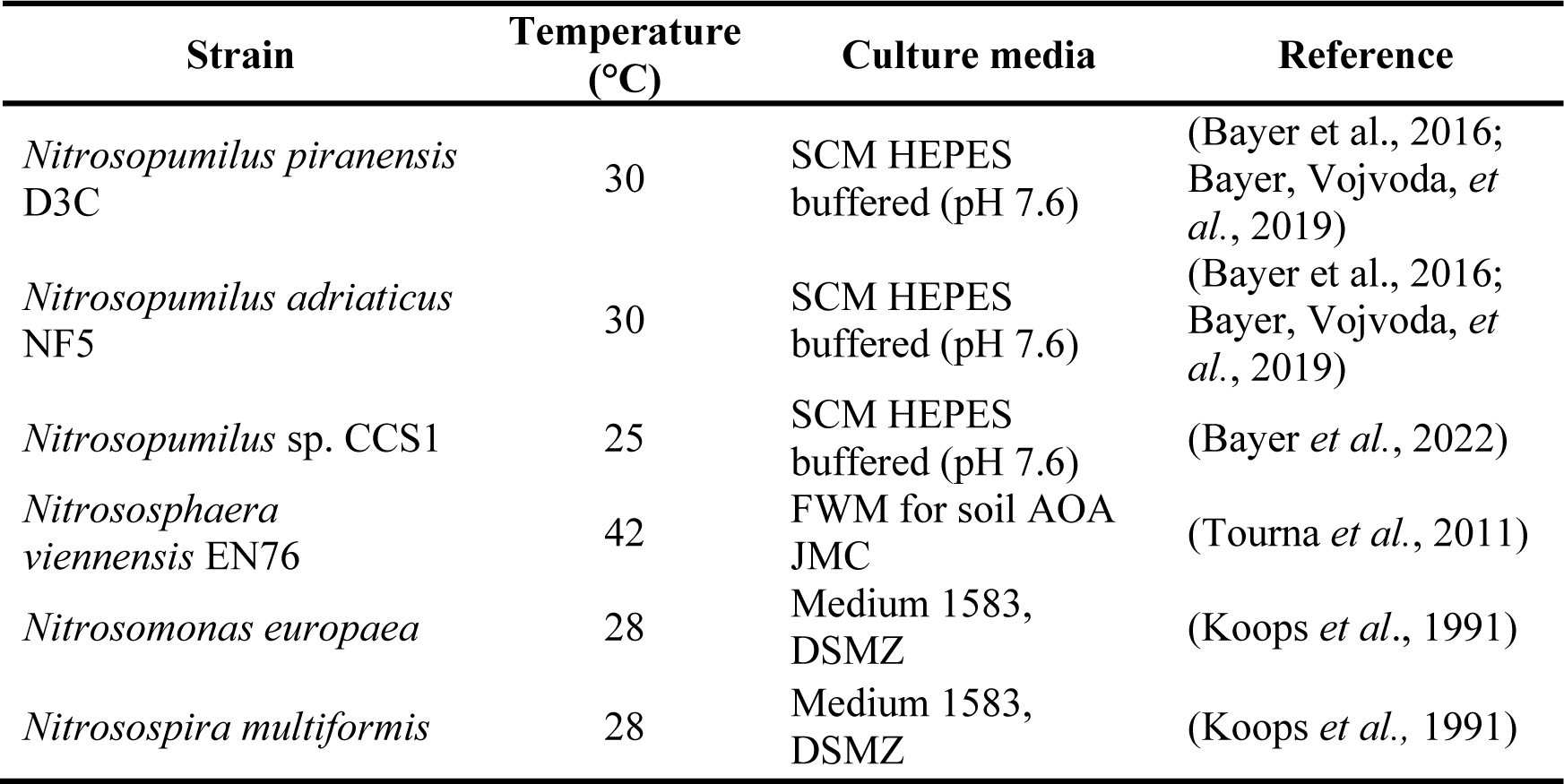
Isolates utilized for batch experiments and culture conditions.

Growth was monitored by ammonium consumption, by nitrite production, and by microscopy. Batch cultures in late exponential phase were used for incubations in oxygen depletion. Immediately before starting the incubations >500µM of NH_4+_ was added to the culture to ensure sufficient ammonia for ammonia oxidation during the experiment.

The batch culture was sparged with argon gas (99.99%) for 45 min to reduce the oxygen concentration in the culture. The culture was sterilely transferred into custom-made glass bottles of 300 ml through a glass tube connection using the overpressure generated in the Ar-sparged culture bottle. The incubation bottles were filled completely without headspace and closed with glass stoppers. The design of the incubation bottles makes oxygen intrusion negligible and includes a long capillary for pressure compensation (Tiano *et al*., 2014). For the incubations, at least three incubation bottles per treatment were used.

Bottles were constantly stirred with glass-coated stirring bars (VWR, UK) at 300 rpm. The bottles were incubated in a water bath at the respective growth temperature of each strain (Table 1), for 72 h. Bottles, glass-coated stirring bars and tube connections were autoclaved before use. Oxygen was monitored constantly during the incubation by trace fluorescence oxygen sensors with a lower detection limit of 0.5nmol·l^−1^ (Lehner et al., 2015), hereafter optodes, that had been glued into the glass bottles prior to utilization. NO concentrations were continuously monitored with NO microsensors (Unisense, Denmark), inserted in the sensor ports of the incubation bottles. NO-microsensors were sterilized with 70% ethanol and rinsed with autoclaved ASTM1a water before insertion into the incubation bottles. NO was previously reported to cause a small (up to 17%) and predictable interference with the optodes. Therefore, all oxygen concentration measurements were corrected for NO interference. Other potential intermediates in ammonia oxidation or nitrite conversion were reported to not interfere with oxygen measurements by the optodes (Kraft *et al.,* 2022).

AOA cultures with a pool of ^15^N-labelled NO_2-_ (1mM) were used to evaluate ^30^N_2_ and ^46^N_2_O production, the product and intermediate of the archaeal NO dismutation (Kraft *et al*., 2022). The ^15^NO_2-_ pool was generated by growing the batch culture with ^15^N labelled NH_4+_ and the culture was used after complete oxidation to ^15^NO_2-_. Oxygen-depleted incubations of AOB strains were not tested with ^15^N labelled substrates as there is existing literature evaluating extensively the formation of ^46^N_2_O from ^15^NO_2-_ during oxygen-depletion (Poth and Focht, 1985; Schmidt *et al*., 2004). Controls with dead cells for a *Nitrosopumilus* strain growing in SCM utilized in the same set up have been already reported in Kraft *et al*. (2022). In the present study, dead controls for cells growing in FWM (see Table 1) were produced by adding 3ml of HgCl_2_ saturated solution in three incubation bottles after oxygen was depleted (Figure S3-B).

Samples were collected at 0, 8, 24, 32, 48, 56 and 72 h with gas-tight syringes (Hamilton, USA) connected to stainless steel needles (Ochs, Germany), through the capillary of the incubation bottles. The volume collected was simultaneously replaced with sterile culture media previously deoxygenated (sparged with Ar) to avoid headspace formation in the incubation bottles. Despite the precautions, a slight intrusion of oxygen when collecting samples could not be avoided.

The samples were preserved in gas-tight 3ml exetainers headspace-free and preserved with 50µl of HgCl_2_ solution. N_2_ and N_2_O were analysed by coupled gas chromatography–isotope ratio mass spectrometry (GC-IRMS) on a Thermo Delta V Plus isotope ratio mass spectrometer (Dalsgaard *et al*., 2012).

## Results

### Oxygen production in AOA

We performed incubations under oxygen depletion for three marine AOA strains from the clade *Nitrosopumilales* (Group I.1a) and one soil AOA strain from the clade *Nitrososphaerales* (Group I.1b) (Figure 1). Prior to the incubations, all the cultures were grown aerobically. Before the start of the incubations, the oxygen concentration in the cultures was reduced by purging with argon and transferred to the incubation bottles, where oxygen consumption started immediately. This immediate oxygen consumption indicated active oxygen respiration and that the cultures were not affected by the purging with argon (Figure 1, first 5 h of incubation).

**Figure 1.**
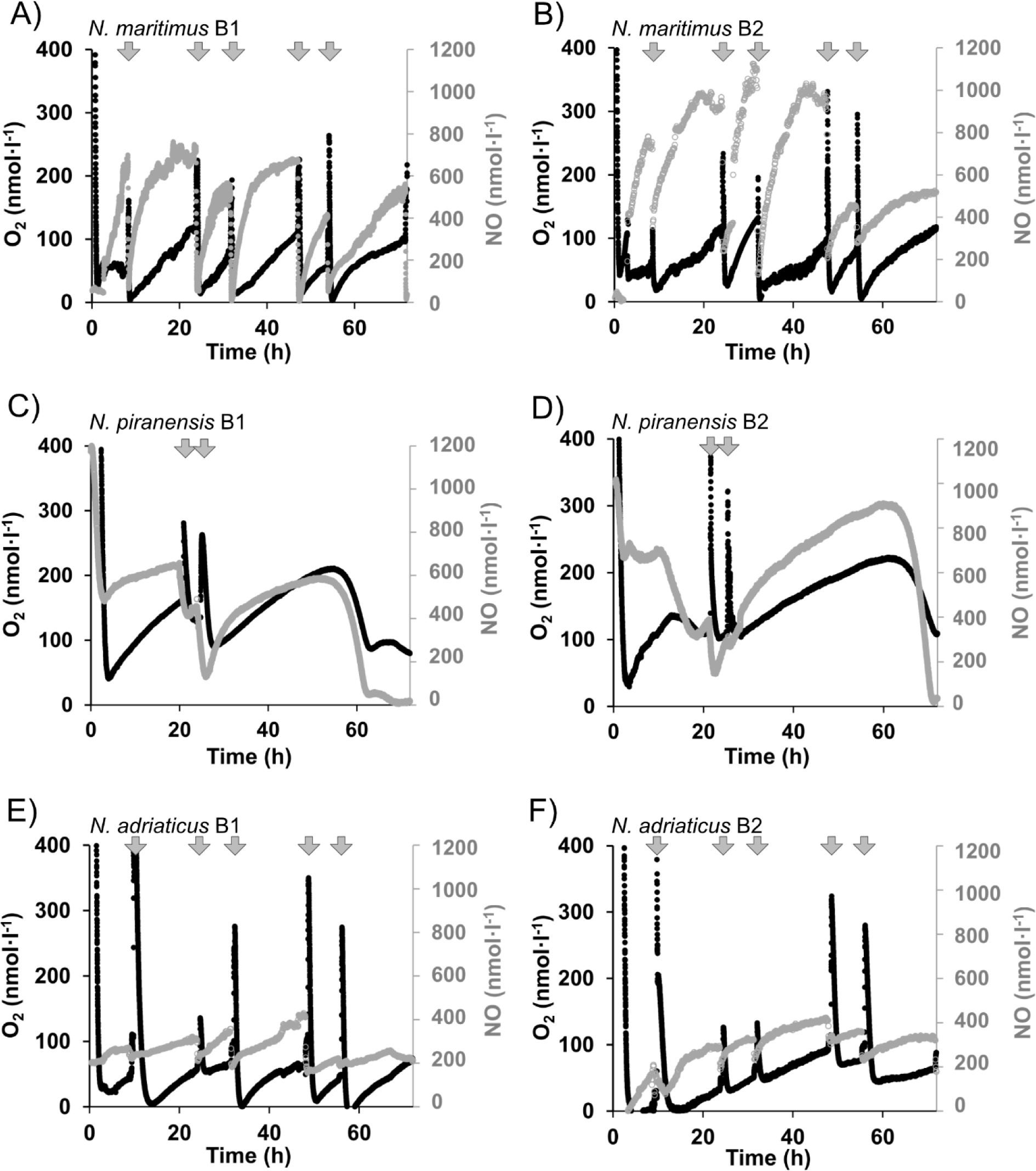
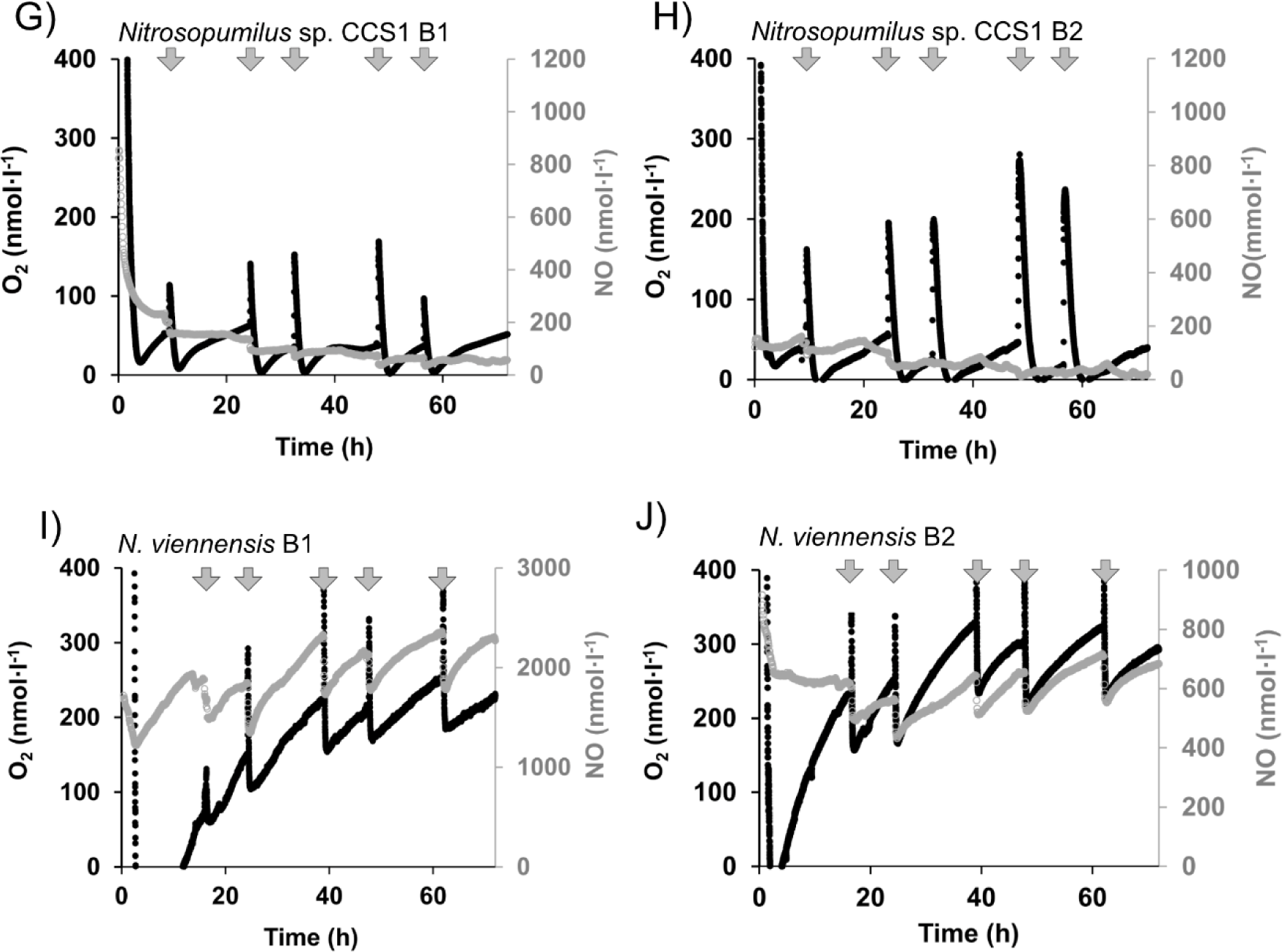
Oxygen accumulation corrected for interference with NO (black symbols) coupled to NO accumulation (grey symbols) in oxygen-depleted incubations of AOA isolates. A, B) *Nitrosopumilus maritimus* SCM1 for which oxygen production was reported before by Kraft *et. al.,* (2022); incubations with the same strain are shown here for comparison. C, D) *Nitrosopumilus piranensis* D3C. E, F) *Nitrosopumilus adriaticus* NF5. Oxygen and NO profile show two panels of reproducible replicates (at least 3n per incubation), the rest of the replicates are showed in the supplementary Figure S2 and S3. Pronounced increases in the oxygen concentration (marked with arrows) are due to oxygen intrusion by sample collection or addition of oxygenated water (see main text). Figure continues in next page. Oxygen accumulation corrected for interference with NO (black symbols) coupled to NO accumulation (grey symbols) in oxygen-depleted incubations of AOA isolates. G, H) *Nitrosopumilus* sp. CCS1. I, J) *Nitrososphaera viennensis* EN76 Oxygen and NO profile show two panels of reproducible replicates (at least 3n per incubation), the rest of the replicates are shown in the supplementary Figure S2 and S3. Pronounced increases in the oxygen concentration (marked with arrows) are due to oxygen intrusion by sample collection or addition of oxygenated water (see maintext).

After oxygen consumption, oxygen accumulation was observed in all the AOA strains tested and it was coupled to NO-accumulation (Figure 1). Abrupt increases in oxygen concentration during incubations, hereafter referred to as oxygen pulses, are a consequence of sample collection or injection of oxygenated water (see material and methods, marked with arrows in Figure 1). Following the oxygen pulses, oxygen was immediately consumed, after which, oxygen accumulation started again, as seen in (Kraft *et al*., 2022).

It is worth noting that the accumulated oxygen (net production) is the result of the balance between the total oxygen produced (gross production) and simultaneous oxygen consumption by cell respiration (including ammonia oxidation), as it has been reported previously for *N. maritimus* SCM1 (Kraft *et al.,* 2022). Oxygen accumulation rates of about 7 nmol·l^−1^·h^−1^ were reached for the two marine strains *N. piranensis* and *Nitrosopumilus* sp. CCS1, while *N. adriaticus* showed a slightly lower accumulation rate (3.9 nmol·l^−1^·h^−1^, Table 2). The soil *N. viennensis* accumulated oxygen at the highest rate among all the observed AOA strains (21 nmol·l^−1^·h^−1^, Table 2).

**Table 2.**
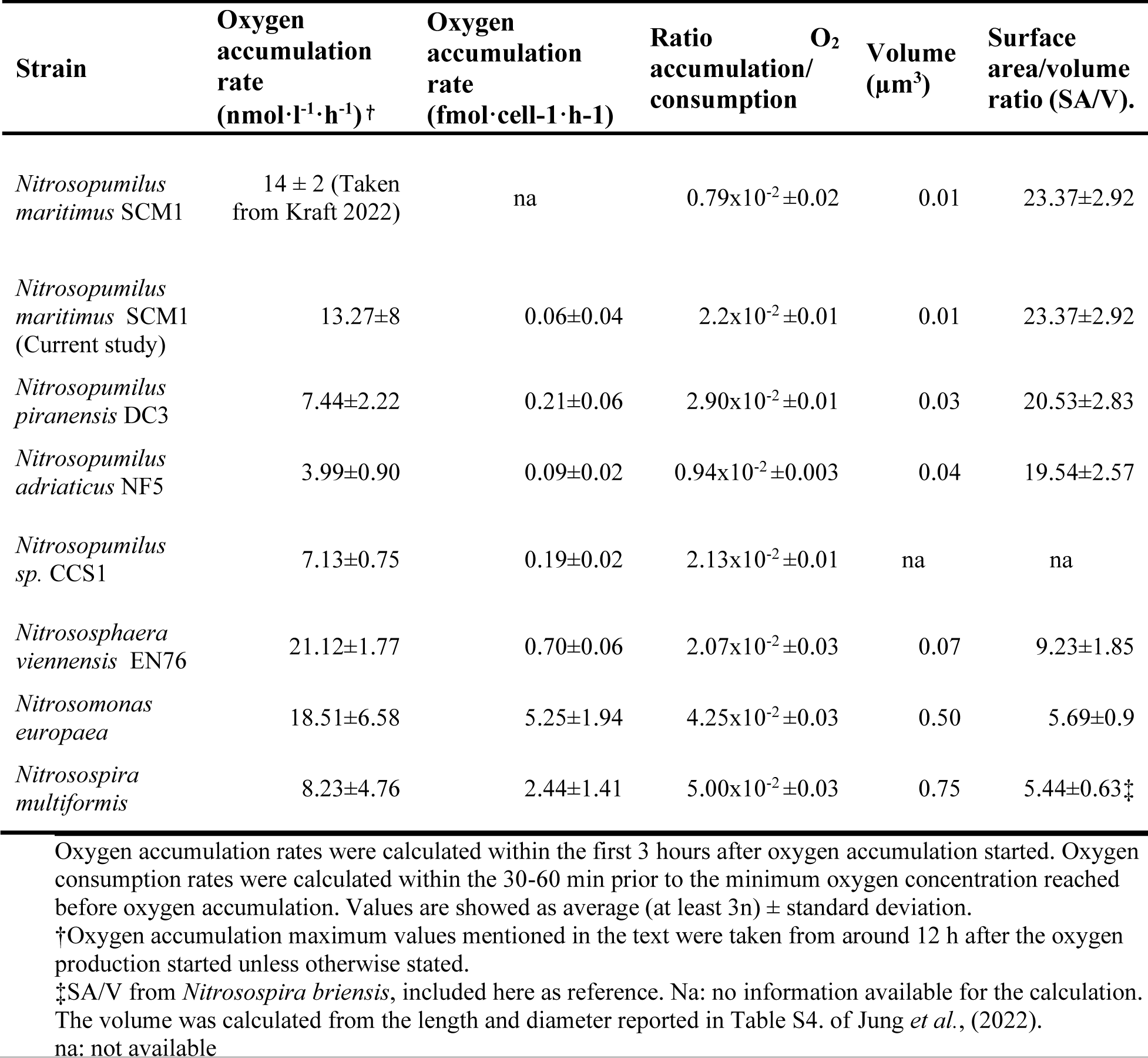
Oxygen accumulation rates in the different AOA and AOB strains and ratio between oxygen respiration rates prior to oxygen production.

The oxygen accumulation rates normalized by cell numbers followed a similar pattern, as the cell densities in all the AOA batch cultures were within the same order of magnitude during the incubation (between 3 to 4·10^4^ cells per ml, except for *N. maritimus* culture which had a cell density of 2·10^5^ cells per ml).

Oxygen accumulated to different levels among the AOA strains, within a range of 50 to 300 nmol·l^−1^. The marine strain *Nitrosopumilus* sp. CCS1 at the lowest end of the range, while the soil strain *N. viennensis* at the highest end. Controls with dead cells did not show any evidence of abiotic oxygen production (Figure S3 B), as was also reported previously by Kraft and collaborators (2022).

Interestingly, the minimum oxygen concentration were oxygen accumulation began varied among strains and, in some cases, among incubation replicates. In most cases, oxygen reached concentrations close to the optode detection limit (0.5 nmol·l^−1^) (Lehner *et al*., 2015), followed by oxygen accumulation, similar to what was observed for *N. maritimus* (Figure 1 A-B) and reported in Kraft *et al.,* 2022. However, *N. piranensis* and *N. viennensis* showed marked differences to this pattern.

In incubations of *N. piranensis*, the lowest oxygen concentrations reached by oxygen respiration, before oxygen accumulation, were always above 38 nmol·l^−1^. In the incubations of *N. viennensis*, the lowest oxygen concentration reached prior to oxygen accumulation was within the first 5h of the incubation. From the second sampling point onwards, after pulses, oxygen was respired, reaching concentrations of between 59 and 260 nmol·l^−1^ (Figure 1 I-J).

In general, the amount of oxygen introduced in the incubation bottles during sample collection, had an influence on the minimum concentration reached before the oxygen accumulation started again, suggesting that a higher oxygen pulse triggered a more pronounced oxygen consumption. For example, in the *N. adriaticus* incubations (Figure 1 E and F), smaller oxygen pulses were followed by less pronounced oxygen consumption, in contrast to the higher oxygen pulses that were followed by a more pronounced oxygen consumption.

The effect of different sizes of oxygen pulses was explored in more detail in incubations with *N. maritimus* SCM1 (Figure S1). The injection of different volumes of sterile oxygenated medium to incubation bottles also showed that higher amounts of oxygen injected were followed by more pronounced oxygen consumption, thus supporting previous observations. While the pattern is the same, the amount of oxygen that triggers complete consumption of oxygen varies among strains.

Oxygen accumulation/consumption ratios were calculated as a proxy for the oxygen production versus consumption activity of the cultures. The ratios among all the AOA tested were similar (around 2·10^−2^), except for the marine strain *N. adriaticus,* which showed the lowest ratio (0.94·10^−2^) (Table 2).

Accumulation of nitric oxide was coupled to O_2_ accumulation in all AOA studied (Figure 1), but with differences in the trends. In the marine AOA strains *N. maritimus*, *N. piranensis* and *N. adriaticus,* as well as in the soil AOA *N. viennensis,* NO accumulation was parallel to oxygen production. In the case of *N. adriaticus*, NO accumulated to a maximum of 600 nmol ·l^−1^, while in *N. maritimus, N. piranensis* and *N. viennensis*, NO concentrations exceeded 600 nmol·l^−1^ and sometimes reached 1000 nmol·l^−1^or more. The lowest NO concentration was accumulated in the incubations with *Nitrosopumilus* sp. CCS1 (up to 200 nmol·l^−1^), moreover, three to four hours after the oxygen accumulation started, NO concentrations declined, while oxygen accumulation continued.

### Oxygen production in AOB

Two AOB strains were incubated under oxygen depletion, following the same methodology described for the AOA strains. Both, *N. europaea* and *N. multiformis* consumed oxygen once the incubation started and reached the oxygen detection limit of the optodes. Surprisingly, after the oxygen was respired, oxygen accumulation was observed. The oxygen accumulation rate of *N. europaea* was twice as high as for *N. multiformis* (Table 2).

When the incubation remained undisturbed (without oxygen pulses), oxygen in *N. europaea* accumulated above 300 nmol·l^−1^ (Figure 2 A and B), which was greater than he other strains tested (AOA and AOB). In the case of *N. multiformis* incubations, oxygen accumulated up to 100 or 200 nmol·l^−1^. Similar to the observations for AOA, after a sample was collected or an oxygen pulse was introduced, oxygen was consumed and oxygen accumulation started again.

**Figure 2.**
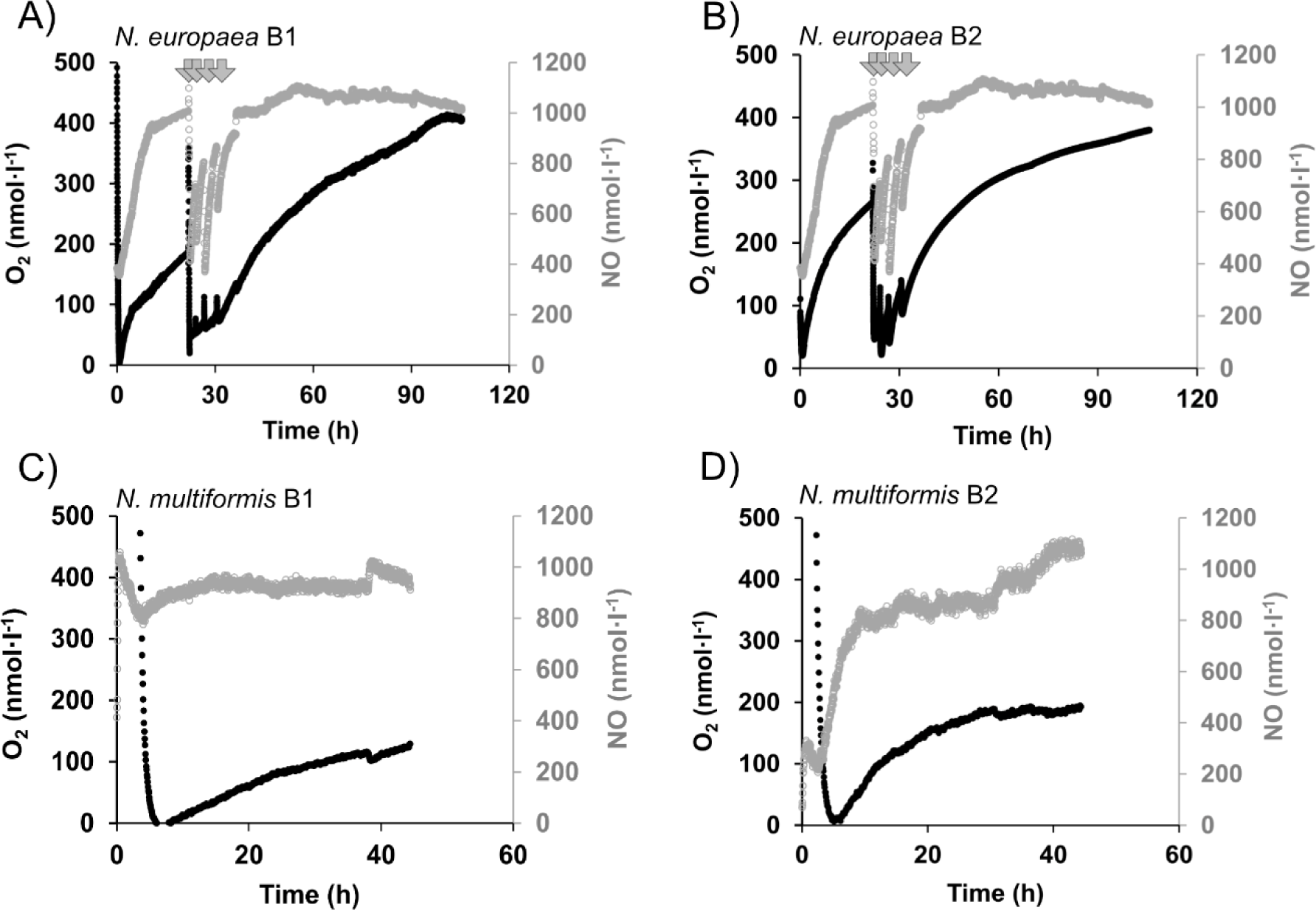
Oxygen accumulation (black symbols) coupled to NO (grey symbols) in oxygen-depleted incubations of different AOB isolates. A, B) *Nitrosomonas europaea* C, D) *Nitrosospira multiformis*. Oxygen and NO profile show two panels of reproducible replicates (at least 3n per incubation). Pronounced increases in the oxygen concentration of A and B are due to oxygen intrusion associated to oxygenated water injection in the incubation (arrows). Oxygen concentrations were corrected for the interference of NO with the optodes.

Cell numbers of AOB cultures were lower than for AOA cultures (around 3×10^3^ AOB cells per ml). Therefore, the oxygen accumulation rates per cell were around 10 to 20 times higher than for AOA. The oxygen accumulation/consumption ratios were similar between the two AOB strains tested, which are notably higher than the calculated ratios for the AOA strains (Table 2).

Just as in the AOA, NO accumulation was coupled to oxygen accumulation in the AOB strains tested (Figure 2). NO concentration in incubations of *N. europaea* increased markedly when oxygen was completely consumed at the beginning of the incubation. After an oxygen pulse, NO concentration decreased, comparable to the observations in the AOA incubations. Interestingly, for both AOB strains, NO reached concentrations around 1000 nmol·l^−1^ and then remained relatively stable.

### N_2_ accumulation and nitrous oxide dynamics under oxygen depletion

N_2_O and N_2_ production from NO_2-_ was tested in oxygen-depleted incubations amended with ^15^NO_2-_ in the marine AOA strains and in the soil AOA (Figure 3). If the AOA strains tested perform NO dismutation as proposed for *N. maritimus,* ^46^N_2_O accumulation and ^30^N_2_ production from ^15^NO_2-_ is expected (Kraft *et al.,* 2022). Production of N_2_O from NO_2-_ by AOB such as *N. europaea* during anoxia, has been explored elsewhere (Poth and Focht, 1985).

**Figure 3.**
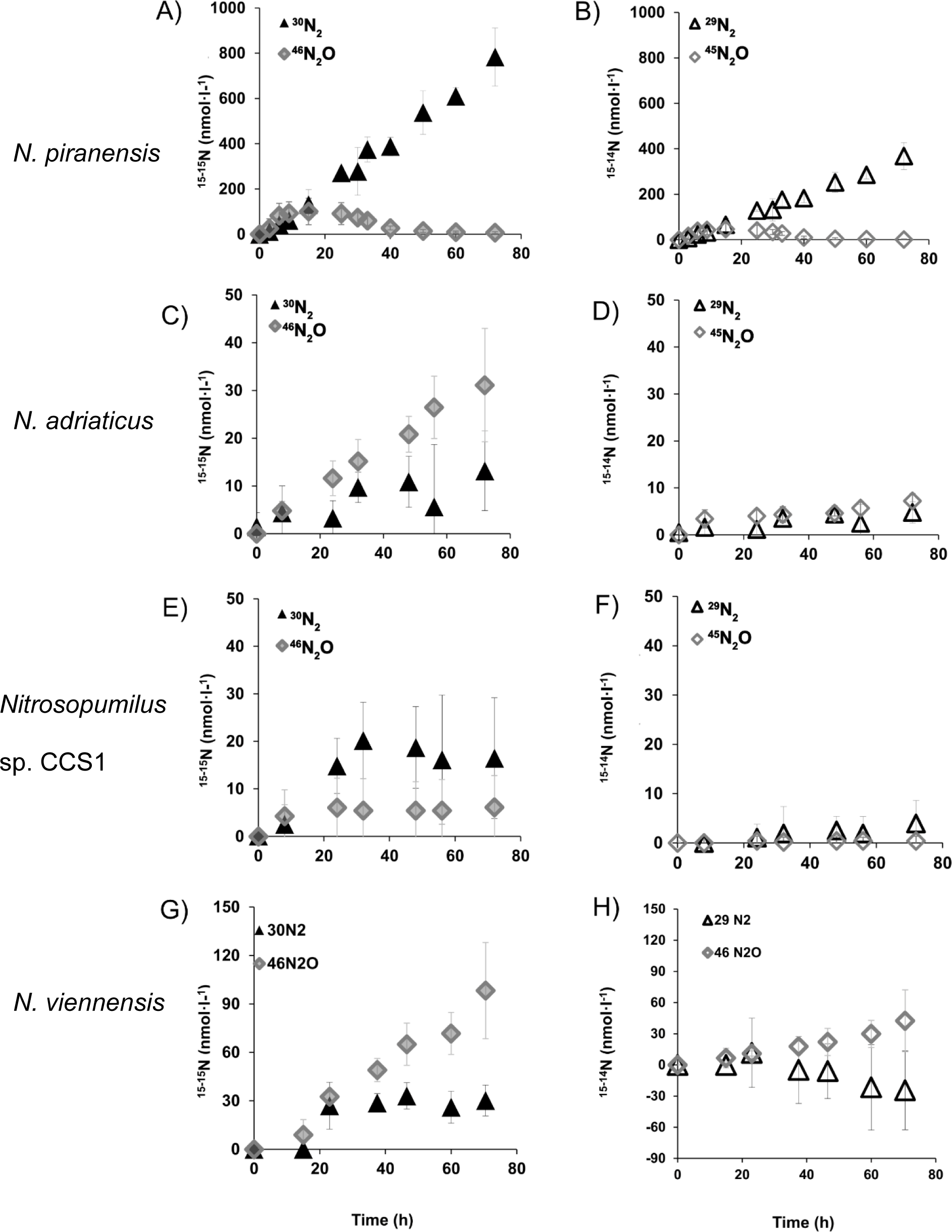
Nitrogen accumulation and nitrous oxide from incubations in oxygen depletion from different AOA isolates with a pool of ^15^NO_2_^−^. Left panels (A, C, E, G) represent ^30^N_2_ production (black triangles) and ^45^N_2_O accumulation (grey diamonds). Right panels (B, D, F, H) represent ^29^N_2_ production (open triangles) and ^44^N_2_O accumulation (open diamonds). Plots in the same row belong to the same strain indicated on the left. Datapoints are average of three replicates and error bars are standard deviation. Some error bars are smaller than the symbols and therefore not visible. For a better interpretation of Figure 3 A-B see the supplementary text.

In the case of the marine AOA *Nitrosopumilus* sp. CCS1 and *N. piranensis,* a transient accumulation of ^46^N_2_O was observed in incubations amended with a pool of ^15^NO_2-_, and ^30^N_2_ accumulated linearly from the beginning of the incubation suggesting that ^15^NO_2-_ is transformed to ^46^N_2_O and then further into ^30^N_2_ (Figure 3 A and E, similar to the observations reported for *N. maritimus* by Kraft and collaborators (2022).

In contrast to the other marine AOA, *N. adriaticus* and the soil strain *N. viennensis* showed a linear increase of ^46^N_2_O from the beginning of the incubation while the simultaneous production of ^30^N_2_ was almost negligible (Figure 3 C and G).

Accumulation of ^29^N_2_ was only observed for the strain *N. piranensis*, for which the incubation was started with ca. 1mM ^15^NO_2-_ and 200µM ^14^NO_2-_ (Figure 3 B; see supplementary text). A slight increase of ^45^N_2_O was observed in the incubation of *N. viennensis* (Figure 3 H).

## Discussion

Oxygen production was observed in all the AOA isolates tested, with representatives of the clades *Nitrosopumilales*, from marine environments, and *Nitrososphaerales* from soils. The coupling of oxygen production to NO accumulation and the transient accumulation of N_2_O followed by N_2_ production (or in some cases only N_2_O production) indicate that oxygen production in these AOA isolates proceeds via the same pathway of NO dismutation as it has been proposed for *N. maritimus* SCM1 (Kraft *et al.,* 2022). These findings make it clear that oxygen production is widely distributed in AOA and therefore have important implications for our understanding of the physiology of AOA in oxygen-depleted ecosystems and consequentially the nitrogen cycle in these environments.

While our results indicate that the core steps of the pathway are the same in the different isolates, we observed interesting differences among them regarding accumulation rates and patterns of oxygen and nitrogen compounds. The soil isolate *N. viennensis* showed a remarkably higher oxygen accumulation rate in comparison to the marine isolates. Considering that the oxygen accumulated (net production) by AOA is a balance between oxygen production and consumption occurring simultaneously (Kraft *et al.,* 2022), the differences in oxygen accumulation rates among the tested strains indicate possible differences in the specific capability of the strains to both, produce oxygen and to utilize it simultaneously.

A noticeable difference among the marine AOA was the minimum oxygen concentration reached after oxygen pulses and prior to oxygen production. This could possibly point towards differences in the oxygen concentration threshold or range that triggers the onset of oxygen accumulation by NO-dismutation in the different AOA.

Aside from variations between isolates, the size of the oxygen pulses determined how much oxygen was consumed before oxygen concentrations increased again. This indicates that the regulation of oxygen production versus consumption, was influenced by the magnitude of the change in oxygen concentrations in the incubation bottles.

In the NO-dismutation pathway proposed for *N. maritimus*, the transient accumulation of ^46^N_2_O and subsequent ^30^N_2_ production from ^15^NO_2-_ was observed (Kraft *et al.,* 2022). The first step of the oxygen-production pathway is the reduction of nitrite to NO and it is most likely performed by the nitrite reductase NirK, which is present in all the AOA strains tested here (Stahl and de la Torre, 2012; Qin *et al*., 2020). The enzymes involved in the NO dismutation and nitrous oxide reduction steps of the proposed pathway are still unknown and neither NO-dismutases nor N_2_O reductases have been identified so far in the available AOA genomes (Kerou *et al*., 2016; Qin *et al*., 2020).

While our results indicate that the core steps of the NO-dismutation pathway are the same in the different isolates, we observed interesting differences in ^46^N_2_O production and ^30^N_2_ accumulation physiology. A striking difference was that *N. adriaticus* and *N. viennensis* produced ^46^N_2_O as the main end product (Figure 3 C and G), in contrast to the other AOA, which showed a transient ^46^N_2_O accumulation together with ^30^N_2_ production. These observations suggest that the enzyme responsible for N_2_O reduction is probably absent or unactive in some AOA.

AOA have been suggested to be important N_2_O contributors in the open ocean, mainly in oxygenated waters and therefore N_2_O production by AOA has mainly been attributed to their aerobic metabolism (Santoro *et al*., 2011; Löscher *et al*., 2012; Wan *et al*., 2023). This study adds evidence for the role of AOA as potential producers of N_2_O from nitrite under oxygen depletion. Most importantly, the difference in N_2_O reduction capability between the AOA tested in the present study, rises the question if certain AOA groups in oxygen-depleted ecosystems can potentially constitute sources of N_2_O to the environment.

A major observation in the present study is the oxygen accumulation by AOB. AOB and AOA perform ammonia oxidation, often occupy different niches mainly due to their different affinities for ammonium and members of both groups can survive in oxygen-limited environments (Martens-Habbena *et al*., 2009; Jung *et al*., 2022). AOB can cope with oxygen-limitation by performing nitrifier denitrification, with some variations among species, such as differences in N_2_O accumulation and N_2_ as final product (Goreau *et al*., 1980; Poth and Focht, 1985; Bock *et al*., 1995). Surprisingly, we observed oxygen production by two distinct and not phylogenetically related AOB strains in pure culture, *N. europaea* and *N. multiformis* when oxygen concentrations reached the low nanomolar range.

On one hand, these results were somehow unexpected, as, despite almost 30 years of research on AOB under oxygen limitation and exposure to anoxia, no oxygen production has been reported to the best of our knowledge. On the other hand, the novelty of our results is based on the use of trace range oxygen sensors. The oxygen accumulation observed in the incubations of the present study reached a maximum concentration of 100 to 300 nmol·l^−1^, while the oxygen sensors previously used had the lowest detection limit in the range of 1 µmol·l^−1^ (Bock *et al*., 1995; Yu *et al*., 2010). In other studies, oxygen concentrations were not explicitly measured, not mentioned (Shrestha *et al*., 2002; Yu *et al*., 2018) or only performed at the beginning of the incubation and/or at the end of the incubation (Cantera and Stein, 2007; Kozlowski *et al*., 2014), leaving the possibility for changes in oxygen concentration during the incubation to be overlooked.

During oxygen depletion, production of ^46^N_2_O from ^15^NO_2-_ by AOB such as *N. europaea* has been reported (Poth and Focht, 1985; Schmidt *et al*., 2004) and identified as nitrifier denitrification. The isotopic signature of the nitrogen compounds is not distinguishable from NO-dismutation.

There are multiple metabolic reactions that can generate N-gases in AOB. For example, *N. europaea* has the capability of producing NO through two mechanisms: aerobic hydroxylamine oxidation to NO by the hydroxylamine oxidoreductase (HAO) and the reduction of NO_2_-to NO by the NirK nitrite reductase. The later one is predominant at low oxygen concentrations (Versantvoort *et al*., 2019). Most of the AOB available in culture have NOR and nitrite reductase (NirK), However *N. europaea* NirK-deficient mutants can still perform nitrifier denitrification (Schmidt *et al*., 2004). Evidence points towards a yet unknown nitrite reductase and holds questions about the function of NirK (reviewed in Stein, 2011).

A study of the transcriptomic response of *N. europaea* to oxygen-limitation, showed that the genes coding for heme-copper-containing cytochrome c oxidases were upregulated during oxygen-limitation. These cytochromes have been suggested to function as a nitric oxide reductase (sNOR) in AOB (Sedlacek *et al*., 2020).

Although there is more information about the physiology of and genes expression in AOB under oxygen depletion than in AOA, no potential NO-dismutase or N_2_O reductase have been identified in AOB, even though N_2_ has been reported as product of the nitrifier denitrification pathway (reviewed in Schmidt, 2008).

Considering the phylogenetic distance between AOB and AOA and concomitantly the differences in their metabolic machineries, the enzyme responsible for oxygen production in AOB may be different from the one in AOA. This would be in agreement with the differences we observed in the NO and O_2_ accumulation patterns.

While AOB occurrence in anoxic environments has been extensively explored, AOA abundance and activity in oxygen depleted ecosystems has been enigmatic and difficult to explain, as the metabolism of these organisms was thought to be strictly aerobic. AOA have been observed in oxygen-depleted marine ecosystems such as oceanic OMZs like the ETNP, ETSP, Arabian Sea and anoxic basins such as the Black Sea (Sinninghe-Damsté *et al*., 2002; Francis *et al*., 2005; Beman *et al*., 2008; Lam *et al*., 2009; Molina *et al*., 2010; Pitcher *et al*., 2011; Bristow *et al*., 2016; Pajares *et al*., 2019; Sollai *et al*., 2019).

Different authors have proposed explanations for AOA abundance in these ecosystems: some suggested the transport from oxic to anoxic depths attached to sinking particles (Ganesh *et al*., 2014), episodic oxygen intrusions in the OMZ that could support aerobic metabolism periodically (Ulloa *et al*., 2012), or even the presence of oxygenic photosynthesis in the upper part of the OMZ, where the oxygen seems to be quickly used by the nitrifier communities (Garcia-Robledo *et al*., 2017). To these feasible explanations, we add the capability to perform an anaerobic metabolism of NO-dismutation as suggested first by (Kraft *et al*., 2022) and expanded in this study.

Here we demonstrated that at least four species of AOA isolated from different oceanic regions produce oxygen. This oxygen production can sustain ammonia oxidation in pure cultures of AOA (Kraft *et al*., 2022). It is still unknown if the same can occur in nature. But if so, the produced oxygen can be used by aerobic members of the surrounding microbial community if their oxygen affinities are sufficiently low (Zakem *et al*., 2020; Canfield and Kraft, 2022).

The discovery of oxygen production upon oxygen depletion by the AOA *N. viennensis*, a representative of an important group of AOA in soil communities (Frey *et al*., 2021; Yin *et al*., 2022), could potentially explain the capability of these nitrifiers to resists oxic-anoxic fluctuations, as a survival mechanism. In terrestrial ecosystems, oxygen depletion can occur when soil remains saturated with water for longer periods. Oxic-anoxic fluctuations can shape the microbial community and cause a change in the nitrifying and denitrifying populations structure and abundances. However, different studies have shown that members of the nitrifiers, including AOA, are tolerant to extended periods of anoxia or can resist oxic-anoxic fluctuations retaining activity after 3-6 weeks (Pett-Ridge *et al*., 2006, 2013).

AOA in soils that perform NO-dismutation might even potentially sustain the metabolic activity of some other members of the community with an extremely low oxygen affinity (nanomolar concentrations), provided that the supply of nitrite would be high enough. An interesting point to further explore is if the production of oxygen is just a survival mechanism that allow the AOA cells to maintain a basal metabolism for a short amount of time or if they can sustain their energy requirements long term via NO-dismutation.

Our results rise the question of the possible competition for resources (nitrite and/or ammonia) between AOA, anammox bacteria and denitrifiers under oxygen depletion. The potential success of AOA performing NO-dismutation in the environment would probably depend on other stressors in the specific environment, that can shape the community and influence the abundance of different microbial cells.

Oxygen production in AOB upon oxygen depletion would have similar environmental implications as discussed previously in the case of AOA, including the potential for ongoing ammonia oxidation in AOB during oxygen depletion and the potential for an unaccounted oxygen source for the surrounding community. Dark oxygen production in two different AOB suggests that, the capability of dark oxygen production from NO is widespread in different microorganisms apart from AOA. Additionally, oxygen production has recently been proposed to occur in *Pseudomonas aeruginosa* (Lichtenberg *et al*., 2021), but there is not isotopic evidence supporting that the mechanism behind is the same as reported for AOA by Kraft *et al*. (2022) and in the present study.

## Conclusions

In conclusion, dark oxygen production is widely distributed among AOA isolates of marine and soil origin.The coupling of oxygen production to NO accumulation and the conversion of nitrite to N_2_O and in some cases subsequently reduced to N_2_, point towards NO-dismutation as the pathway, with some physiological differences among the AOA strains. One of the most striking differences was the lack of N_2_O reduction capability by *N. adriaticus* and *N. viennensis,* leading to N_2_O accumulation as final product, different from the rest of strains tested, which produced N_2_. This difference challenges our understanding about the role of the AOA as potential sources or sinks of the greenhouse gas N_2_O in oxygen-depleted ecosystems. Furthermore, we presented evidence of oxygen production in two strains of ammonia oxidizing bacteria upon oxygen depletion, with higher oxygen production rates in comparison to AOA. It remains to be investigated wether dark oxygen production by AOA and AOB occurs in the environment and if the oxygen produced can be utilized by other community members in nature.

## Acknowledgements

This study was funded by the Villum Foundation grant no. 00025491 to Beate Kraft. We thank Martin Könneke for providing the strain *N. maritimus* SCM1 and Alyson Santoro for providing the strain *Nitrosopumilus* sp. CCS1 and *N. adriaticus* NF5 as well as her valuable contributions to this study. We also thank Bo Thamdrup and Laura Bristow for their support and advice when processing samples on the IRMS.

## Data Availability Statement

All data generated or analysed during this study are included in this published article and its supplementary information files.

