## supplementary_text_S1_S2_S3 for "Oxygen production via NO dismutation in different ammonia oxidizers"

[1]Nordcee. Department of Biology, Faculty of Sciences. University of Southern Denmark.

[†]Corresponding authors: Beate Kraft, Nordcee. Department of Biology, Faculty of Sciences. University of Southern Denmark. Campusvej 55, DK-5230 Odense M

Elisa Hernández-Magaña, Nordcee. Department of Biology, Faculty of Sciences. University of Southern Denmark. Campusvej 55, DK-5230 Odense M

**Supplementary text.**

Prior to the incubation of *N. piranensis* with $^{15}NO_2^-$ 1L (Figure 3 A-B), the batch culture was aerobically growth with $^{14}NH_4^+$, and when it reached late exponential phase, the volume of the culture was upscaled to 5L with 1mM of $^{15}NH_4^+$ and used for the incubation when reached late exponential phase was again reached. Therefore, the culture started with a pool of ca. 1mM $^{15}N$-$NO_2$- and 200µM $^{14}N$-$NO_2^-$.

### Supplementary figures.

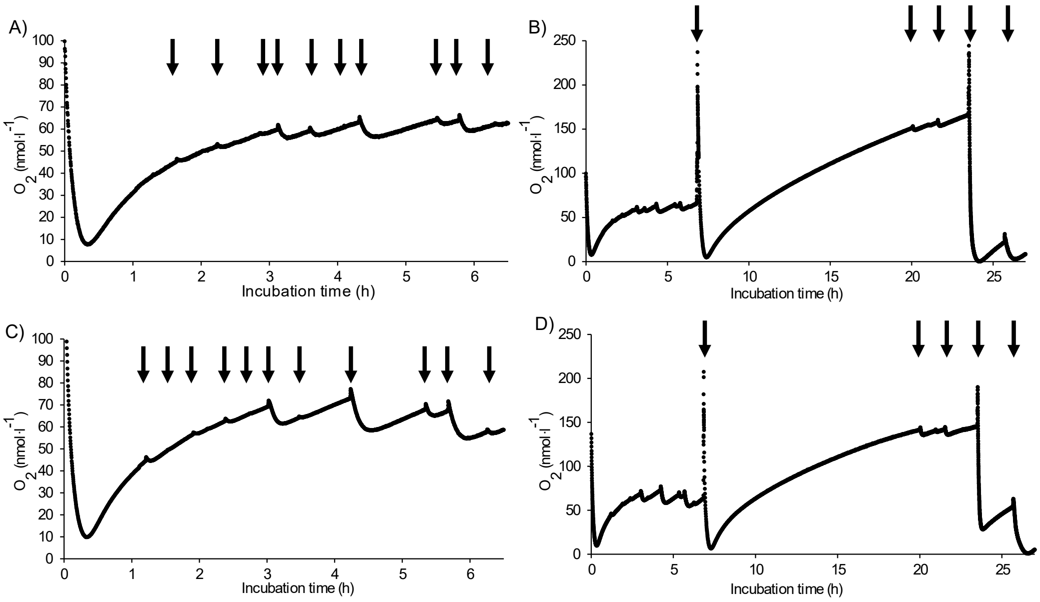


**Figure S.1.** Response of oxygen producting incubations of *N. maritimus* SCM1 to different sizes of oxygen pulses. Two replicates (A, B and C, D) are shown. A and C show the first 6h of the incubations, while B and D give an overview of the whole incubation time. Arrows indicate oxygen pulses.

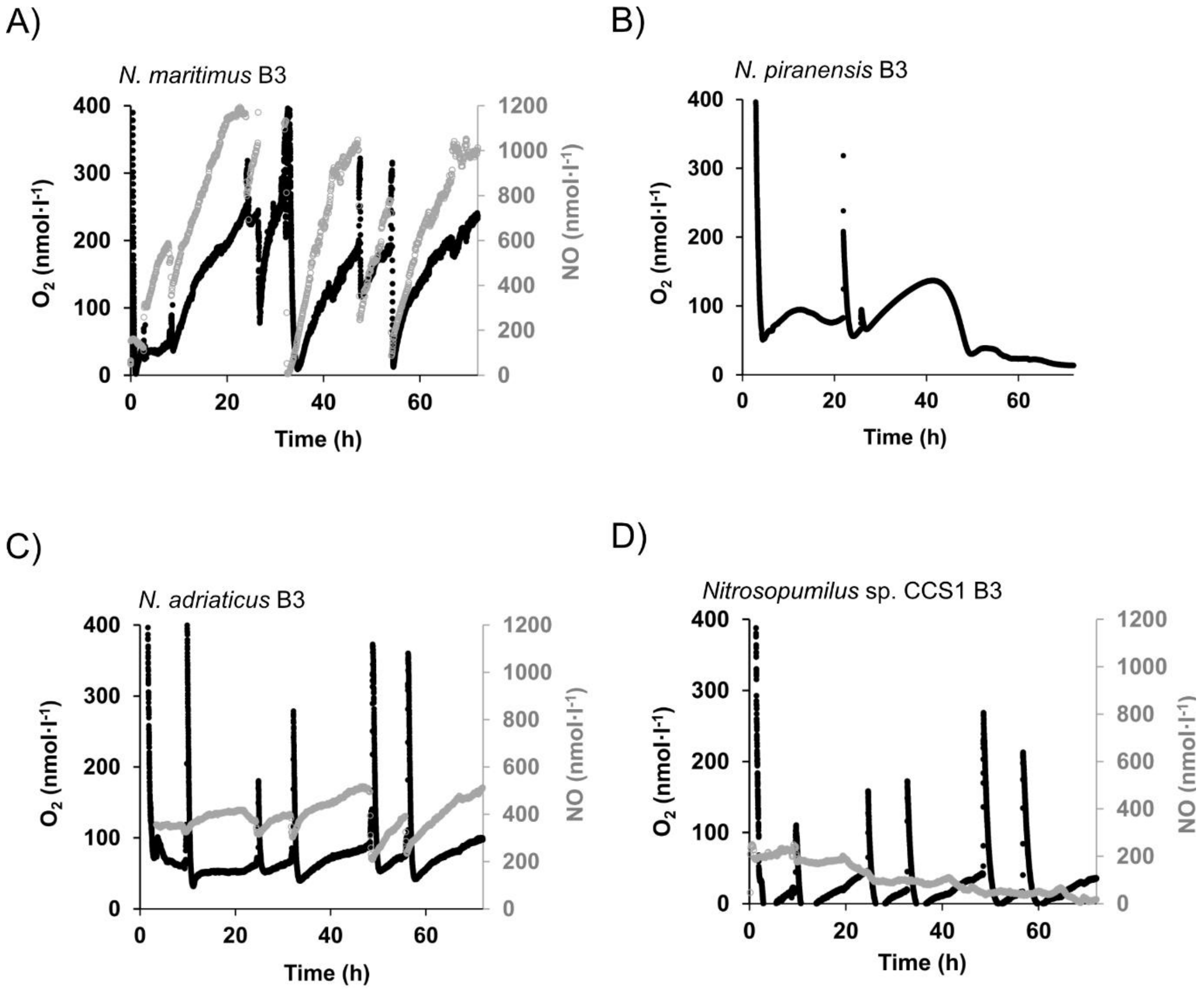


**Figure. S2.** Oxygen accumulation (black symbols) coupled to NO (grey symbols) in oxygen-depleted incubations of different AOA isolates. Oxygen and NO profile show one panel of reproducible replicates (at least 3n per incubation), the rest of the replicates are showed in the main figure 1. Pronounced increases in the oxygen concentration are due to oxygen intrusion associated to sample collection. Oxygen concentrations were corrected for the interference of NO with the optodes. Incubation of *Nitrosopumilus piranensis* B) did not have NO measurement of that specific replicate.

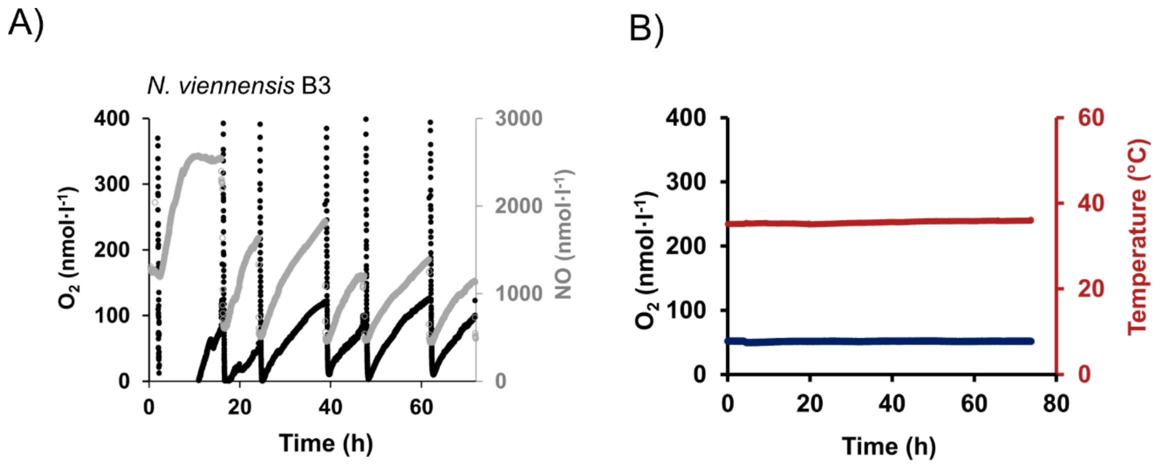


**Figure S.3.** A) Oxygen accumulation (black symbols) coupled to NO (grey symbols) in oxygen-depleted incubations of AOA from soil *Nitrosophaera viennensis*. Oxygen and NO profile show one panel of reproducible replicates (at least 3n per incubation), the rest of the replicates are showed in the main figure 1. Pronounced increases in the oxygen concentration are due to oxygen intrusion associated to sample collection Oxygen concentrations were corrected for the interference of NO with the optodes. B) Killed control with medium and dead cells of *N. viennensis* with concentrated mercuric chloride. Incubated parallel to replicates of B1-B3 presented on Figure 1 and Figure S. 3. A. Blue line are oxygen concentrations, red indicates the temperature at what the incubation was stablished. No oxygen consumption or production was observed from the media components.
